# Angiotensin II disrupts the cytoskeletal architecture of human urine-derived podocytes and results in activation of the renin-angiotensin system

**DOI:** 10.1101/2021.03.18.436037

**Authors:** Lars Erichsen, Martina Bohndorf, Md. Shaifur Rahman, Wasco Wruck, James Adjaye

## Abstract

High blood pressure is one of the major public health problems which causes severe disorders in several tissues including the human kidney. One of the most important signaling pathways associated with the regulation of blood pressure is the renin-angiotensin system (RAS), with its main mediator angiotensin II (ANGII). Elevated levels of circulating and intracellular ANGII and aldosterone lead to pro-fibrotic, -inflammatory and -hypertrophic milieu that causes remodelling and dysfunction in cardiovascular and renal tissues. Furthermore, ANGII has been recognized as major risk factor for the induction of apoptosis in podocytes, ultimately leading to chronic kidney disease (CDK).

In the past, disease modeling of kidney-associated malignancies was extremely difficult, as the derivation of kidney originated cells is very challenging. Here we describe a differentiation protocol for reproducible differentiation of SIX2-positive urine derived renal progenitor cells (UdRPCs) into mature podocytes bearing typical foot processes. The UdRPCs-derived podocytes show the ability to execute Albumin endocytosis and the activation of the renin-angiotensin system by being responsive to ANGII stimulation. Our data reveals the ANGII dependent downregulation of *NPHS1* and *SYNPO*, resulting in the disruption of the complex podocyte cytoskeletal architecture, as shown by immunofluorescence-based detection of α–ACTININ. In the present manuscript we confirm and propose UdRPCs as a unique cell type useful for studying nephrogenesis and associated diseases. Furthermore, the responsiveness of UdRPCs-derived podocytes to ANGII implies potential applications in nephrotoxicity studies and drug screening.

## Introduction

The kidney glomerulus or renal corpuscle consists of a glomerular tuft and the Bowman’s capsule. Its major task is the filtration of blood to generate urine. The glomerulus consists of distinct cell types: endothelial cells, mesangial cells, parietal epithelial cells of Bowman’s capsule and podocytes, which are attached to the outer part of the glomerular basement membrane (GBM). Podocytes are pericyte-like cells with a complex cellular organization. Key characteristics are the cell body with major and minor foot processes (FPs) [1]. The FPs contain a highly organized actin-based cytoskeleton, essential for maintaining the complex architecture typical of podocytes. Alpha-actinin-4 (ACTN4) and Synaptopodin (SYNPO) are both highly expressed in podocyte foot processes and function as cross-linkers of F-actin filaments in order to bundle them and thereby enhancing podocyte signaling and mobility [2, 3]. Filtration slits are formed by the spatial arrangement of FPs of neighboring podocytes and each of these slits is bridged by the so called glomerular slit diaphragm (SD), which is a comparable structure to the adherens junctions [1]. The most abundant proteins that contribute to SD formation are Nephrin (NPHS1) [4] and Podocin (NPHS2) [5]. Furthermore, Nephrin is also associated with the actin cytoskeleton thereby contributing to podocyte actin dynamics and FPs formation [6]. Together the FPs and SD establish the filtration barrier of the kidney with its selective permeability.

High blood pressure is one of the major public health problems, causing severe disorders in several tissues including heart, brain and kidney [7]. One of the most important signaling pathways in blood pressure regulation is the renin-angiotensin system (RAS), with its main mediator angiotensin II (ANGII). At the molecular level, ANGII signaling is mediated by two classes of receptors (AGTR1 and AGTR2). Both receptors are expressed in a wide variety of tissues (including the heart, kidney, blood vessels, adrenal glands and cardiovascular control centers in the brain) and upon stimulation control vasoconstriction [8, 9]. Human Podocytes express both types of ANGII receptors and are indeed effector cells for this peptide [10]. Furthermore, elevated levels of ANGII have been identified as a main risk factor for the initiation and progression of chronic kidney disease (CKD). Increased ANGII concentrations are associated with the downregulation of Nephrin and Synaptopodin expression in podocytes [11, 12]. The depletion of NPHS1 and SNYPO is causative for podocyte injury [13], which is typically associated with marked albuminuria [1], and increased podocyte apoptosis [14]. This is especially crucial since mature podocytes are terminal differentiated cells that are unable to undergo cell division *in vivo* [1]. As a result of this, the replenishment capabilities of podocytes are limited and expose the glomerulus vulnerable to exogenous noxae. Long lasting hazardous cues, such as high blood pressure, can manifest in the significant loss of podocytes, which is a hallmark in the development of CKD [15].

In the past disease modeling for kidney-associated malignancies was extremely difficult, as the derivation of kidney originated cells is very challenging. This is particularly true for podocytes, since their complex architecture is not well preserved from kidney biopsy tissue [16]. To model podocyte-related diseases, researchers either used an immortalized podocyte cell line [16–18] or iPSC-based models [19–23]. Whilst the first approach shows the typical drawback of immortalized cells, namely chromosomal aneuploidies, thus making these cells prone to cancerous transformation. The iPSC-based differentiation approach lacks reproducibility and attainment of mature podocytes with elaborate foot processes.

We recently reported human urine as a non-invasive source of renal stem cells with regenerative potential [24]. The urine derived renal progenitor cells (UdRPCs) express renal stem cell markers such as SIX2, CITED1 WT1, CD133, CD24 and CD106. Stimulation of UdRPCs with the GSK3ß-inhibitor (CHIR99021) induced differentiation into renal epithelial proximal tubular cells. In the present study, we provide a detailed protocol for the direct differentiation of SIX2-positive UdRPCs into mature podocytes with typical foot processes without the need of immortalized cells or pluripotent stem cells. We provide the full characterization of the generated podocytes at the transcriptome, secretome and cellular level. Furthermore, with ANGII treatment we demonstrate the responsiveness of the podocytes to the renin-angiotensin system (RAS) with a downregulated expression of NPHS1 and SYNPO.

Our data demonstrates and establishes human urine-derived podocytes as a valuable *in vitro* model for studying podocyte injury, loss and ultimately CKD. In addition, we hypothesize that the cells can also be used for further drug testing and eventually kidney-associated regenerative therapies.

## Material and Methods

### Cell culture conditions

UdRPCs were cultured in Proliferation Medium (PM) composed of 50% DMEM high Glucose and 50% Keratinocyte medium supplemented with 5% FCS, 0.5% NEAA, 0.25% Gtx and 0.5% Penicillin and Streptomycin at 37°C (Gibco, Carlsbad, United States) under hypoxic conditions. For differentiation, the cells were seeded at low and high density in 6-or 12-well plates coated with 0,2% type 1 Collagen (Gibco) and grown in PM for 24 hours. After this the medium was changed to advanced RPMI (Gibco) supplemented with 0.5% FCS, 1% Penicillin and Streptomycin and 30 µM retinoic acid. Typical podocyte morphology was observed after 7 days. ANG II (Sigma Aldrich, St. Louis, Missouri, United States) was diluted in the culture medium to a final concentration of 100 µM. Cells were incubated for 6h and 24h with ANGII and the conditioned medium was kept for secretome analyses.

### Relative Quantification of podocyte associated gene expression by real-time PCR

Real time PCR of podocyte associated gene expression was performed as follows:

Real time measurements were carried out on the Step One Plus Real Time PCR Systems using MicroAmp Fast optical 384 Well Reaction Plate and Power Sybr Green PCR Master Mix (Applied Biosystems, Foster City, United States). The amplification conditions were denaturation at 95°C for 13 min. followed by 37 cycles of 95°C for 50s, 60°C for 45s and 72°C for 30s. Primer sequences are listed in supplement table 1.

### Bisulfite genomic sequencing

Bisulfite sequencing was performed following bisulfite conversion with the EpiTec Kit (Qiagen, Hilden, Germany) as described [25]. Primers were designed after excluding pseudogenes or other closely related genomic sequences which could interfere with specific amplification by amplicon and primer sequences comparison in BLAT sequence data base (https://genome.ucsc.edu/FAQ/FAQblat.html). PCR primers are listed in supplementary table 2.

Briefly, the amplification conditions were denaturation at 95°C for 13min. followed by 37 cycles of 95°C for 50s, 54°C for 45s and 72°C for 30s. The amplification product is 270 bp in size. Amplification products were cloned into pCR2.1vector using the TA Cloning Kit (Invitrogen, Carlsbad, United States) according to the manufacturer’s instructions. On average 30 clones were sequenced using the BigDye Terminator Cycle Sequencing Kit (Applied Biosystems) on a DNA analyzer 3700 (Applied Biosystems) with M13 primer to obtain a representative methylation profile of each sample. 5’-regulatory gene sequences are referring to +1 transcription start of the following sequence:

Homo sapiens WT1 transcription factor (WT1), RefSeqGene (LRG_525) on chromosome 11

NCBI Reference Sequence: NG_009272.1

Homo sapiens NPHS2, podocin (NPHS2), RefSeqGene (LRG_887) on chromosome 1

NCBI Reference Sequence: NG_007535.1

### Immunofluorescence staining

Cells were fixed with 4% paraformaldehyde (PFA) (Polysciences, Warrington, United States). To block unspecific binding sites the fixed cells were incubated with blocking buffer containing 10% normal goat or donkey serum, 1% BSA, 0.5% Triton, and 0.05% Tween, for 2h at room temperature. Incubation of the primary antibody was performed at 4°C overnight in staining buffer (blocking buffer diluted 1:1 with PBS). After at least 16h of incubation the cells were washed three times with PBS/0.05% Tween and incubated with a 1:500 dilution of secondary antibodies dilution. Afterwards the cells were washed again three times with PBS/0.05% Tween and nuclei were stained with Hoechst 1:5000 (Thermo Fisher Scientific, Waltham, United States) and podocyte cytoskeleton was stained with Alexa Flour 488 phalloidin (Thermo Fisher Scientific) (1:400). Images were captured using a fluorescence microscope (LSM700; Zeiss, Oberkochen, Germany) with Zenblue software (Zeiss). Individual channel images were processed and merged with Fiji. Detailed Information of the used antibodies are given in supplementary table 3.

### Microarray data analyses

Total RNA (1 μg) preparations were hybridized on the PrimeView Human Gene Expression Array (Affymetrix, Thermo Fisher Scientific, USA) at the core facility Biomedizinisches Forschungszentrum (BMFZ) of the Heinrich Heine University Düsseldorf. The raw data was imported into the R/Bioconductor environment and further processed with the package affy using background-correction, logarithmic (base 2) transformation and normalization with the Robust Multi-array Average (RMA) method. The heatmap.2 function from the gplots package was applied for cluster analysis and to generate heatmaps using Pearson correlation as similarity measure. Gene expression was detected using a detection-p-value threshold of 0.05. Differential gene expression was determined via the p-value from the limma package which was adjusted for false discovery rate using the q value package. Thresholds of 1.33 and 0.75 were used for up-/down-regulation of ratios and 0.05 for p-values. Venn diagrams were generated with the Venn Diagram package. Subsets from the venn diagrams were used for follow-up GO and pathway analyses as described by Zhou et al [26]. Gene expression data will be available online at the National Centre of Biotechnology Information (NCBI) Gene Expression Omnibus.

### Secretome analyses

Conditioned media from control podocytes as well as ANG II-treated cells were analysed using the Proteome Profiler Human Kidney Biomarker Array Kit distributed by Research And Diagnostic Systems, Inc. (Minneapolis, Minnesota, United States) as described by the manufacturer. Obtained images were analyzed by using the Image J software [27] with the Microarray Profile plugin by Bob Dougherty and Wayne Rasband (https://www.optinav.info/MicroArray_Profile.html). The integrated density generated by the Microarray profile plugin function *Measure RT* was used for follow-up processing which was performed in the R/Bioconductor environment [28]. Arrays were normalized employing the Robust Spline Normalization from the Bioconductor lumi package [29]. A threshold for background intensities was defined at 5% of the range between maximum and minimum intensity and a detection-p-value was calculated according to the method described in Graffmann et al. [30].

### Cluster analysis of the Renin-angiotensin pathway

Pathways and associated genes were downloaded from the KEGG database [31] in July 2020 and annotated with official gene symbols. Genes from the KEGG pathway hsa04614 Renin-angiotensin system were extracted from the microarray data normalized with the RMA method from the package oligo [32] in R/Biocondcutor [28] as described above and subjected to the R *heatmap()* function using Pearson correlation as similarity measure and color scaling per row.

### Albumin endocytosis assay

The functionality of podocyte differentiated from urine-derived renal progenitor cells was analyzed employing the Albumin endocytosis assay. In brief, podocytes were plated at low density of 10 % in a 12-well plate coated with 0,2% type 1 collagen. 48 hrs later the cells were washed with 1x PBS and replaced with podocyte culture medium supplemented with 20 μg/ml of BSA-Alexa Fluor™ 488 and incubated at 37°C for 60 minutes. After incubation, the cells were washed 3X with cold PBS and fixed with 4% PFA for 15 minutes. Finally, the images were taken at the excitation wavelength of 488 nm and an emission wavelength of 540 nm using a LSM700 florescence microscope.

## Results

### Generation of human podocytes from urine derived renal progenitor cells

We recently reported a “*rice grain*” fibroblast-like morphology resembling MSCs isolated directly from urine samples of 4 male (UM) and 6 female (UF) donors. These cells express the renal stem cell markers SIX2, CITED1 WT1, CD133, CD24 and CD106, we referred to these cells as urine derived renal progenitor cells (UdRPCs) [24]. UdRPCs maintain their stem cell features for almost 12 passages (fig. 1a). So far, we have established a differentiation protocol leading to renal epithelial proximal tubular lineage by supplementation with the GSK3ß-inhibitor (CHIR99021) or to podocytes as described in fig. 1a. UdRPCs acquire typical podocyte “*fried egg*” shape morphology after 7 days, when cultured at 70% confluency in adv. RPMI medium supplemented with retinoic acid. This protocol was applied to cells isolated from human urine of several individuals (age ranged between 45 and 76 years), resulting in the robust generation of podocytes (fig. 1a). For further analysis we evaluated mRNA expression of the two most abundant proteins within the FPs and SD of human podocytes, Nephrin (*NPHS1*) and Synaptopodin (*SYNPO*) respectively by quantitative real time PCR. We analysed mRNA expression of podocytes derived from three individuals, two males (UM48 and UM51) and a female (UF45) and compared it to their undifferentiated counterparts. The expression of *NPHS1* and *SYNPO* was upregulated in the differentiated podocytes compared to the respective UdRPCs (fig. 1b+c). To show methylation changes within the 5’-regulatory region of the key podocyte transcription factor WT1, we applied bisulfite genomic sequencing. In total a 270 bp long WT1 promoter fragment spanning 19 CpG-dinucleotides was analysed. We found, that the undifferentiated UdRPCs UM51 had a dense methylation pattern at CpG position 12 and 13 of the 5’-regulatory region (fig. 1d). Upon applying our differentiation protocol, methylated DNA at the respective positions for UM51-derived podocytes, was found to be almost completely lost (fig. 1e). As a mark of renal cell functionality, Albumin endocytosis was observed after exogenous BSA was supplemented to the culture medium (fig. 1f).

**Figure 1:**
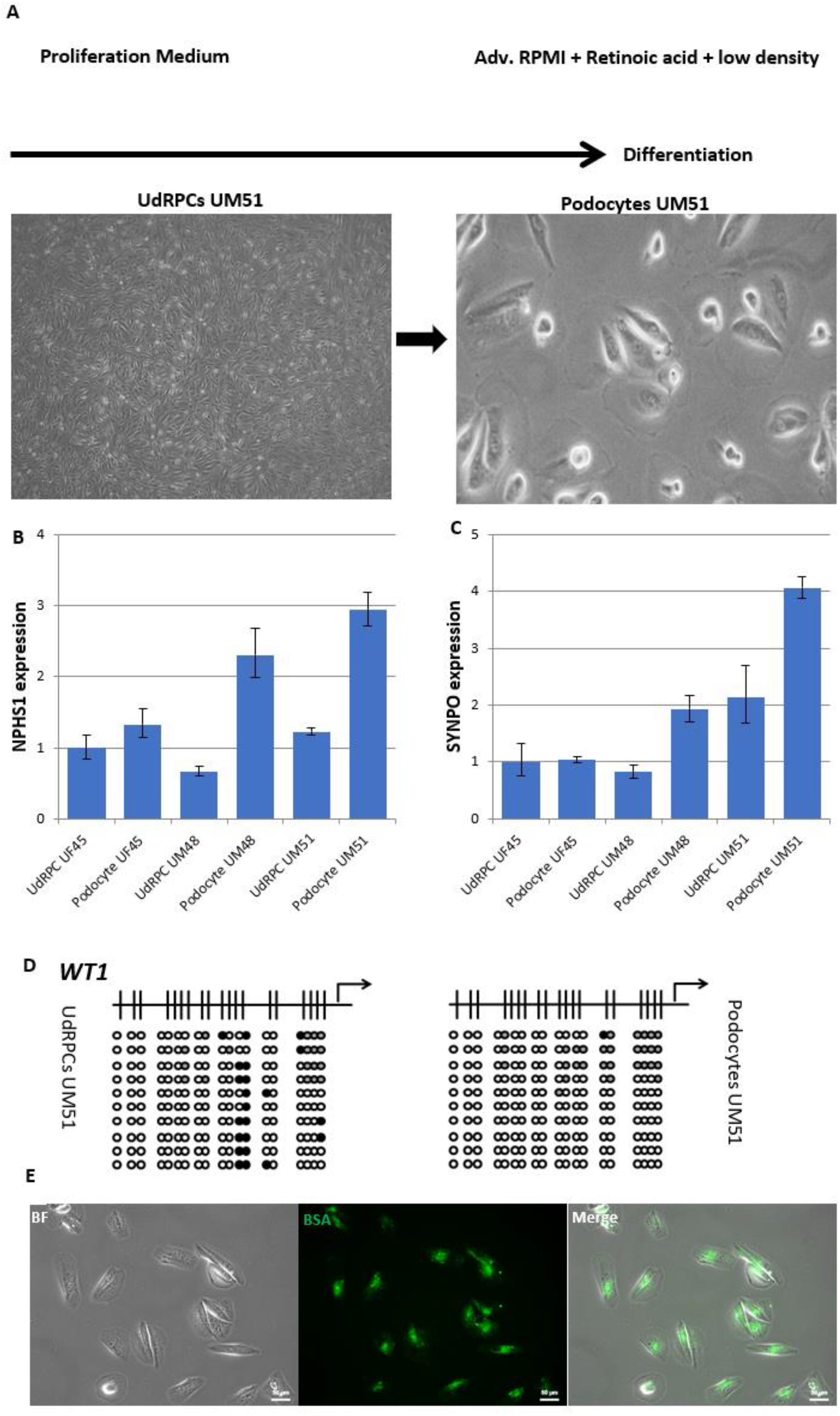
Derivation of mature and functional podocytes from human urine derived renal progenitor cells. The self-renewal capacity of UdRPCs is maintained by proliferation medium. Differentiation of UdRPCs into podocytes is induced by low density cultivation in adv. RPMI supplemented with 30µM retinoic acid (a). Expression of the podocyte associated genes *NPHS1* (b) and *SYNPO* (c) was determined by quantitative real time PCR. Bisulfite genomic sequencing of a 270bp long WT1 promoter fragment, spanning 19 CpG-dinucleotides, provide detailed information about the dynamic DNA Methylation changes occurring during the differentiation process into podocytes (d). Black, white and grey circles refer to methylated, unmethylated and undefined CpG dinucleotides, respectively. UdRPCs show dense methylation patterns at CpG position 12 and 13, whilst differentiated podocytes show a lack of methylation. Infiltration of exogenous BSA supplemented into the culture medium confirmed endocytosis of Albumin in UM51 podocytes (e).

### Retinoic acid enhances podocyte differentiation

To improve the differentiation protocol of UdRPCs into podocytes, we supplemented the adv. RPMI medium with 30 µM retinoic acid and analysed the derived UM51-podocytes by immunofluorescence-based detection of expression of Nephrin (NPHS1), Podocin (NPHS2) and CD2AP (fig. 2a-f). While the UM51 podocytes cultured in adv. RPMI already show expression of all three proteins, which are necessary for podocyte functionality, the addition of retinoic acid significantly increased the protein expression levels of these key podocyte markers. Additionally, we measured the expression of *LMX1b, NPHS1, SYNPO* and *WT1* by quantitative real time PCR (fig. 2g-j). While the addition of retinoic acid had only minor impact on the expression of *LMX1b* and *WT1*, the expression levels of *NPHS1* and *SYNPO* were highly increased, by 3 and 2-fold, respectively.

**Figure 2:**
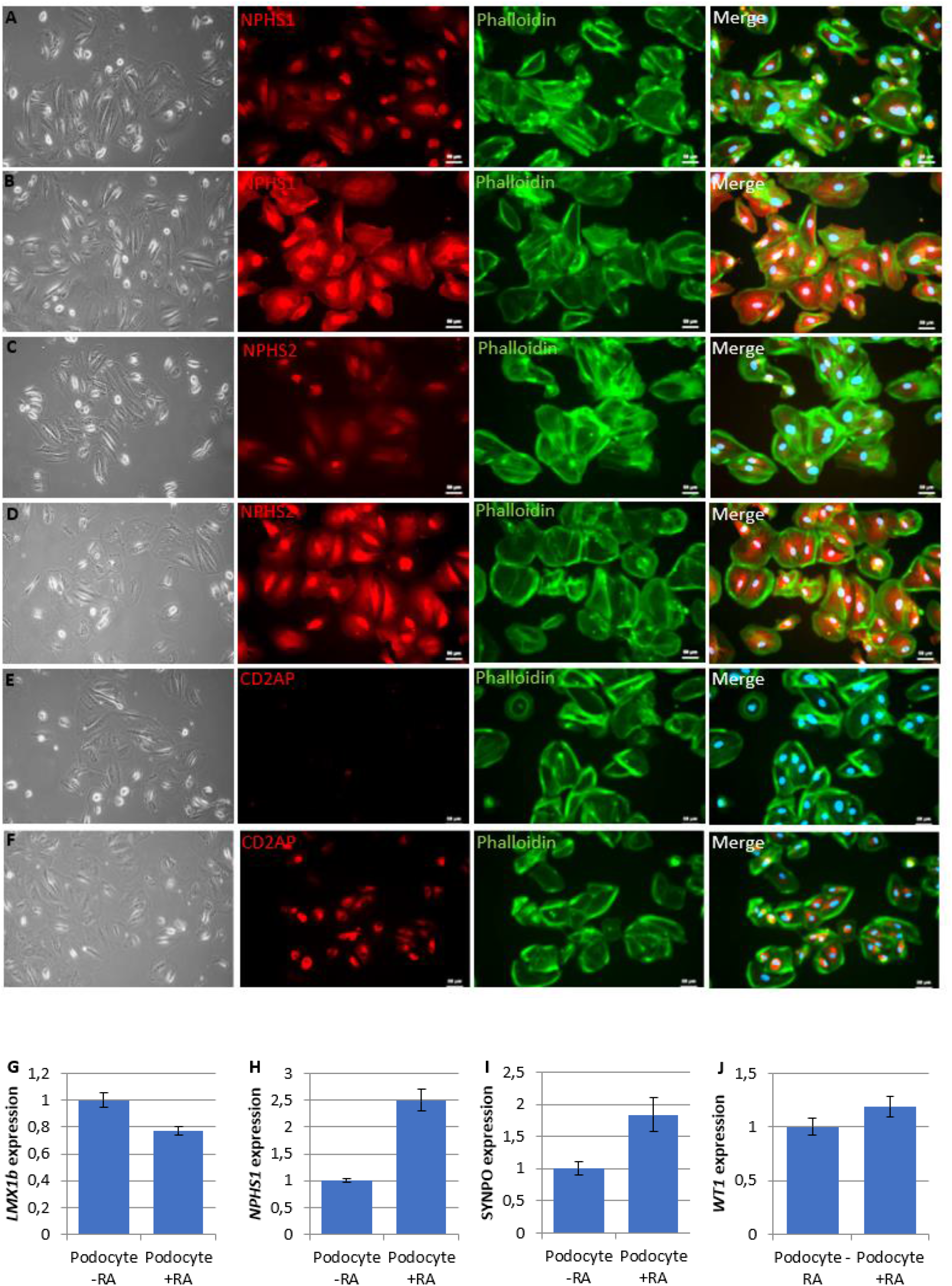
Retinoic acid enhances the maturation of UdRPC-derived podocytes. UdRPCs differentiated into podocytes, by low density cultivation in advanced RPMI medium supplemented without (a, c, e) and with (b, d, f) 30 µM retinoic acid. Immunofluorescence-based detection of NPHS1 (a, b), NPHS2 (c, d) and CD2AP (e, f) expression. Podocyte cytoskeleton was stained with phalloidin. Expression of podocyte markers *LMX1b* (g), *NPHS1* (h), *SYNPO* (i), and *WT1* (j) was determined by quantitative real time PCR.

### Comparative transcriptome analysis of Urine derived Renal Progenitor cells with their podocyte counterparts

After the successful and reproducible differentiation of UdRPCs into podocytes we performed a comparative transcriptome analysis of UF45, UM48 and UM51. Hierarchical clustering analysis comparing the transcriptomes of UdRPCs with podocytes revealed a distinct expression pattern of both cell types (fig. 3a). By comparing the expressed genes (det-p < 0.05) 250 are exclusively expressed in the UdRPCs and 600 in podocytes (fig. 3b). The most over-represented GO BP-terms exclusive to UdRPCs are associated with cell division and activation of the pre-replicative complex. In comparison the most over-represented GO BP-terms exclusive to podocytes are associated with cell fate commitment, cell morphogenesis and regulation of ion transport (fig. 3c). These genes include important regulators of podocyte morphology and function such as: *BCAS3, BDNF, CLIC5, CXCR4, DNM3, GPER1, GRIA3, LCN2, LPAR3, MBL2, NFASC, NRXN1, ROBO1, SH3GL2, SYT1, WNT1, WNT8A, WNT9A*, and the collagens *COL12A1* and *COL14A1*. In total, 13762 genes were found to be expressed in common between UdRPCs and podocytes, of these 671 were upregulated and 904 downregulated in podocytes (fig. 3b). The downregulated genes are again associated with cell cycle processes and methylation (fig. 3e). Interestingly genes associated with methylation include *DNMT3A, EZH1, MTR, SUZ12, TET2* and genes associated with the cell cycle *BUB1, BUB1B, CDC7, CDC25A, FGFR2, MAD2L1, MAP2K6, SIX3, TOP2A, ZBTB16*. In contrast, upregulated genes are associated with cell adhesion, positive regulation of cell migration, cell substrate adhesion, morphogenesis of an epithelium and regulation of cytokine production (fig. 3d). These include *ARHGEF2, CD44, CXCL16, CX3CL1, DPP4, ETS1, ITGA2, ITGA6, ITGAV, LAMA3, LAMB3, LAMC1, LAMC2, NOTCH1, PDGFA, PDGFB, PODXL, SEMA3A, SEMA3C, WNT7A* and *VEGFC*. Furthermore, genes annotated under the GO BP-term morphogenesis of an epithelium are known to be critical effectors either for epithelial-mesenchymal transition (EMT) or mesenchymal-epithelial transition (MET) *ALDH1A3, CAV1, DLG1, FZD7, MICAL2, PDGFA, SMAD3, SMURF1, SMURF2, TGM2, TNFAIP3* and *VEGFC*. We suggest that our differentiation protocol trigger these processes and thereby mimic a process that is referred to as branching morphogenesis during mammalian kidney development [33]. The generated podocytes express numerous genes which are annotated with neuronal GO BP-terms, such as axonogenesis, axon guidance, chemotaxis, and regulation of axonogenesis (supplementary table 4). We hypothesize that this might reflect the migration process during podocyte maturation or a necessary task during the formation of the foot processes. Additionally, the transcriptomes of UdRPCs UF45, UM48 and UM51, their derived podocytes were compared with expressed genes in iPS cell–derived podocytes, kidney biopsy isolated human glomeruli, and mouse podocytes (supplementary fig. 3), as reported by Sharmin et al. [21]. Furthermore, we analysed the expression of several Solute Carrier (SLC) Family members (supplementary fig. 3). Both analyses revealed distinct expression patterns of UdRPCs and their differentiated counterparts. In particular, the SLC genes were found to be exclusively upregulated in podocytes. Key genes associated with the complex podocyte morphology, like *PODXL, NPHS1, NPHS2* and *SYNPO* were found to be upregulated in the differentiated podocytes of all three individuals. Interestingly undifferentiated UdRPCs express *CD2AP* (an important stabilizer of the slit diaphragm), which seem to be unaffected by our differentiation protocol. In addition, our results provide detailed information about genes exclusively expressed in human UdRPCs which are mesenchymal and differentiated podocyte which are epithelial (supplementary tables 4, 5 and 6).

**Figure 3:**
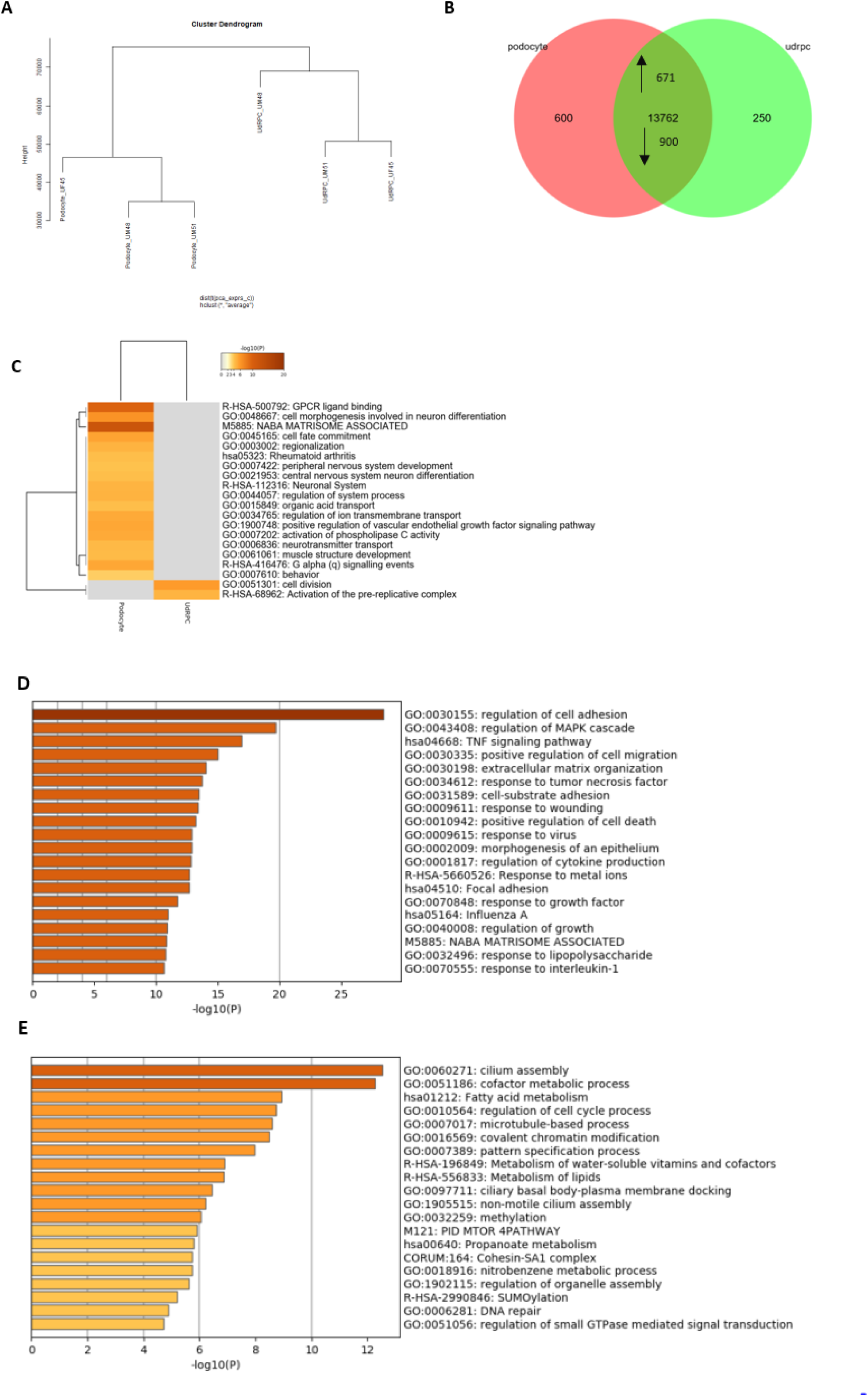
Comparative transcriptome and Gene Ontology analysis of urine-derived renal progenitors UF45, UM 48, UM51 and derived podocytes. A hierarchical cluster dendrogram revealed distinct clusters of UdRPCs and their derived podocytes. (A). Expressed genes (det-p < 0.05) in UdRPCs and podocytes compared in the Venn diagram (B), shows distinct (600 in podocytes; 200 in UdRPCs) and overlapping (13762) gene expression patterns. Of the overlapping genes, 671 are upregulated and 900 downregulated in podocytes. The most over-represented GO BP-terms exclusive in either UdRPCs or podocytes are shown in c and including cell division and activation of the pre replicative complex for the UdRPCs and cell fate commitment, cell morphogenesis and regulation of ion transport for the podocytes. The upregulated 671 genes in podocytes, are associated with cell adhesion, positive regulation of cell migration, cell substrate adhesion, morphogenesis of an epithelium and regulation of cytokine production, while downregulated 900 genes are associated with cell cycle processes and methylation (E).

### Effect of Angiotensin II (ANGII) on podocyte morphology and expression of podocyte-specific genes

Elevated levels of ANGII have been identified as a main risk factor for the initiation and progression of chronic kidney disease (CKD), furthermore increased ANGII concentrations are associated with the downregulation of Nephrin and Synaptopodin expression in podocytes [11, 12]. To evaluate the capacity of our generated podocytes modelling acute and chronic kidney injury, we prepared a final concentration of 100 µM ANGII in adv. RPMI medium supplemented with 30 µM retinoic acid and treated podocytes derived from three individuals, two-males African (UM48 and UM51) and one female-Caucasian (UF45), for 6h and 24h respectively. This analysis was performed for podocytes cultured under low (fig. 4) as well as high cell density (supplementary fig. 1). After 6h, dynamic changes in morphology could be observed by immunofluorescence-based detection of α –ACTININ expression (fig. 4a + supplementary fig. 1a). While untreated podocytes showed the typical “fried egg” morphology, ANG II treated podocytes derived from all three individuals underwent massive disruption of the cytoskeleton, resulting in the inhibition of podocyte spreading and subsequent loss of foot processes as observed by the round and condensed morphology (fig. 4a supplementary fig. 1a). To confirm that the disruptive effect is indeed mediated by ANGII, the effects on the cytoskeleton were evaluated by gene-specific mRNA expression of both ANGII receptors, *AGTR1* and *AGTR2*, as well as podocyte key structural proteins *NPHS1* and *SNYPO* and the pro-inflammatory cytokine IL-6 (fig. 4b-f). While *AGTR1* expression was found not to be significantly downregulated, *AGTR2* was found to be upregulated in podocytes derived under low density cultivation from all three individuals after 6h (fig. 4b-c). Interestingly when the cells were cultured under high density conditions both receptors seemed to be upregulated due to the ANGII treatment after 6h, with the only exception for *AGTR2* in UF45 podocytes (supplementary fig. 1b-c). When the ANGII treatment was prolonged to 24h, the mRNA expression level of *AGTR1* was still downregulated in UM48 podocytes cultured under low density conditions, at a similar level as in 6h, while UM51 podocytes show a modest increase 0.6-fold. mRNA expression of *NPHS1* and *SYNPO* was downregulated upon ANGII treatment in all individuals and culture conditions, except for the *SYNPO* expression in podocytes UF45 under high density cultivation which was upregulated (fig. 4d-e and supplementary fig. 1d-e). Podocytes cultured under low density conditions showed only minor changes in the *NPHS1* and *SYNPO* expression after 6h of ANGII treatment, while mRNA expression in UM51 and UM48 podocytes cultured under high density was found to be downregulated by 2.6- and 1.7-fold (*NPHS1*) and 2.6- and 1-fold (*SYNPO*) respectively (supplementary fig. 1b-c). When ANGII treatment was prolonged to 24h under low density conditions, UM48 and UM51 podocytes both showed downregulated expression of *NPHS1* expression, 0.9-fold, and 3-fold, respectively. In contrast expression of *SYNPO* remained at similar levels seen in 6h of ANGII treatment. Since ANGII is a pro-inflammatory factor we also evaluated mRNA expression of *IL-6* in UF45, UM48 and UM51 podocytes after 6h and UM48 and UM51 after 24h of ANGII treatment (fig. 4f). *IL-6* mRNA levels were found not to be significantly regulated after 6h (UF45: 0.9-fold; UM48: 0.3-fold and UM51: 0.3-fold) or 24h of ANGII treatment (UM48: 0.3-fold and UM51: 0.5-fold). To evaluate whether the transcriptional changes of cytoskeleton-associated genes could be in part mediated by epigenetic modifications, we applied bisulfite genomic sequencing to analyse the methylation status of a CpG-rich region of the *NPHS2* promoter. In total a 415bp long *NPHS2* fragment 11 bp upstream of the TSS, spanning 23 CpG-dinucleotides, was analyzed and this revealed a lack of methylation changes occurring during 6h of ANGII treatment in UM51 podocyte (supplementary fig. 2).

**Figure 4:**
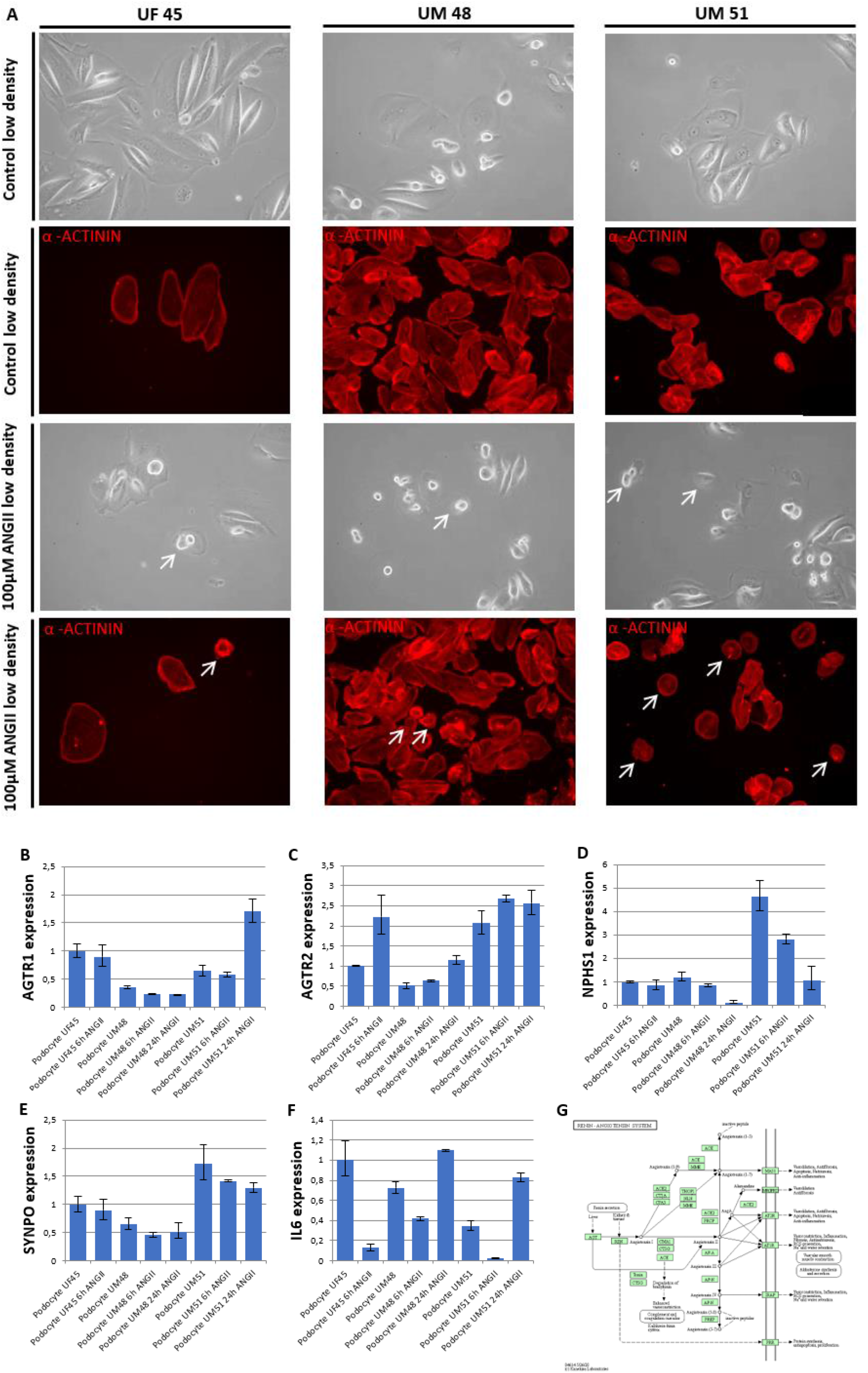
Dynamic changes in the morphology of human urine derived podocytes post treatment with Angiotensin II (ANGII) after 6h and 24h. UdRPC-differentiated podocytes from UF45, UM48 and UM51. The top panel (phase contrast) shows the typical “fried egg” shaped podocyte morphology. The lower two panels show morphology changes after 6h of 100 µM ANGII treatment. Podocyte cytoskeleton was visualized by immunofluorescence-based detection of α –ACTININ in red (a). ANG II interferes with the cytoskeleton of the podocytes, inhibiting podocyte spreading and resulting in the loss of foot processes and the observed roundish phenotype (indicated by the white arrows). Expression of ANGII receptors *AGTR1* (b), *AGTR2* (c), expression podocyte markers *NPHS1* (d), *SYNPO* (e) and pro inflammatory cytokine *IL-6* (f) were determined by quantitative real time PCR normalized with the ribosomal encoding gene-RPL0. An overview of the KEGG annotated renin-angiotensin system is given in figure 4g.

### Effects of Angiotensin II on the transcriptome and secretome of urine derived podocytes

To further investigate the effect of ANGII, we performed a comparative transcriptome analysis of UM48 and UM51 podocytes after 6h and 24h of ANGII treatment (n=2 biological replicates per genotype). Hierarchical clustering analysis comparing the transcriptomes of non-treated and treated podocytes revealed distinct transcriptomes for the three conditions, without treatment, 6h and 24h of ANGII treatment respectively (fig. 5a). Interestingly while podocytes treated for 6h seem to cluster rather by their genetic background, the cells treated for 24h formed a distinct cluster. Further analyses identified 478 genes (det-p < 0.05) exclusively expressed in the control conditions and 63 genes in the 6h Angiotensin conditions (fig. 5b). The most over-represented GO BP-terms derived from the 478 genes are associated with growth and locomotion (fig. 5d). These genes include *BDNF, DNAH5, TTC21A* and *VPS13A*. In contrast the most over-represented GO BP-terms linked to the 63 are associated with immune system process, response to stimulus and cell proliferation (fig. 5d). These genes include *GATA2, GLI3, IL12RB1, LILRB2, PECAM-1, SIX3* and *VTCN1*. In total 13783 non-regulated genes were found to be expressed in common between both conditions. By comparing the expressed genes (det-p < 0.05) after 24h of ANGII treatment, 339 genes were found to be exclusively expressed in the control podocytes and 335 genes in the treated cells (fig. 5c). The most over-represented GO BP-terms of the 339 genes are associated with multi-organism and multicellular organismal process and locomotion (fig. 5e). These genes include *ASPM, CAMK2B, HTR6, ITPR1, ISL1, ISL2, KCNMB4, NCAPH, ROBO1, SCN3B, SHANK3, SLIT1, TOPA2* and *TYRO3*. The full gene list can be found in supplementary table 7. In total 13922 genes were found to be expressed in common between both conditions, from which 1287 were upregulated and 1100 downregulated after 24h of ANGII treatment (supplementary tables 8+9). The upregulated genes are associated with cell cycle, DNA repair, cell cycle phase transition, methylation, and regulation of cellular response to stress (fig. 5f). These include *ALMS1, ATF4, CDK6, DDIT3, EDN1, H2AC4, H2BC14, H3C7, FANCD2, FOXM1, MSH2, MSH6, NUP35, NUP37, NUP54, NUP107, NUP133, NUP153, NUP155, NUP214, PARP2, PCNA, PLK2, PRC1, RAD21, RAD51C, SNAI1, SNAI2, SMC2, SMC4, SUZ12, TGFB2, WDR70, WNT5A* and *XRCC4*. Interestingly genes annotated for the GO BP-terms cell cycle and methylation were also found in the undifferentiated UdRPCs. The downregulated genes after 24h ANGII treatment are associated with regulation of cell adhesion, cell junction organization, axonogenesis, actin cytoskeleton organization, tissue-, and gland morphogenesis (fig. 5g). Genes annotated under the GO BP-term regulation of cell adhesion include *AFDN, CD74, CORO1A, CORO2B, DDR1, DPP4, EPCAM, ITGA2, JAG1, KLF4, LAMA3, LAMA5, LDLR, NLRP3* and *PODXL*. Genes annotated under the GO BP-term cell junction organization include *COL17A1, CLDN3, LAMB3* and *LAMC2*. Genes annotated under the GO BP-term tissue and gland morphogenesis include *CXCL16, ELF3, NTN4, PAX6, PLAU, SEMA3A, SLC9A3R1, SMURF1, TUBB2B, VANGL2, WNT7A, WNT7B* and *WNT10A*. Of note, expression of *CXCL16, DPP4, ITAG2, LAMA3, LAM3B, PODXL, ROBO1, SEMA3A*, SMURF1 and WNT7A were found to be upregulated upon differentiation of UdRPCs into podocytes, but then downregulated upon ANGII treatment. This finding may suggest that these genes might have major implications during the differentiation of UdRPCs into podocytes and ANGII induced podocyte injury. Finally, expression of numerous genes annotated under the GO BP-term actin cytoskeleton organization were downregulated upon 24h ANGII treatment and revealed a distinct clustering of UdRPCs, Podocytes and Podocytes treated with ANGII for 24h. The full gene list is given in supplementary tables 4+9. Interesting to note most of these genes also show basal levels of expression in undifferentiated UdRPCs. Genes annotated for the GO BP-term actin cytoskeleton organization include *ANKRD13D, ANKRD27, COL5A2, KRT17, KRT19, NRCAM, SEMA3A, SEMA3F, SEMA4C, SEMA5A, SEMA6D, SEMA7A, SLC9A3R1, SLC12A7, SLC12A8, TICAM1* and *VANGL2*. These findings further emphasize the disruptive effect of ANGII on cytoskeleton of podocytes as it has been observed by immunofluorescence-based detection of protein expression and q-RT PCR. An overview of the KEGG annotated renin-angiotensin system is given in figure 4g.

**Figure 5:**
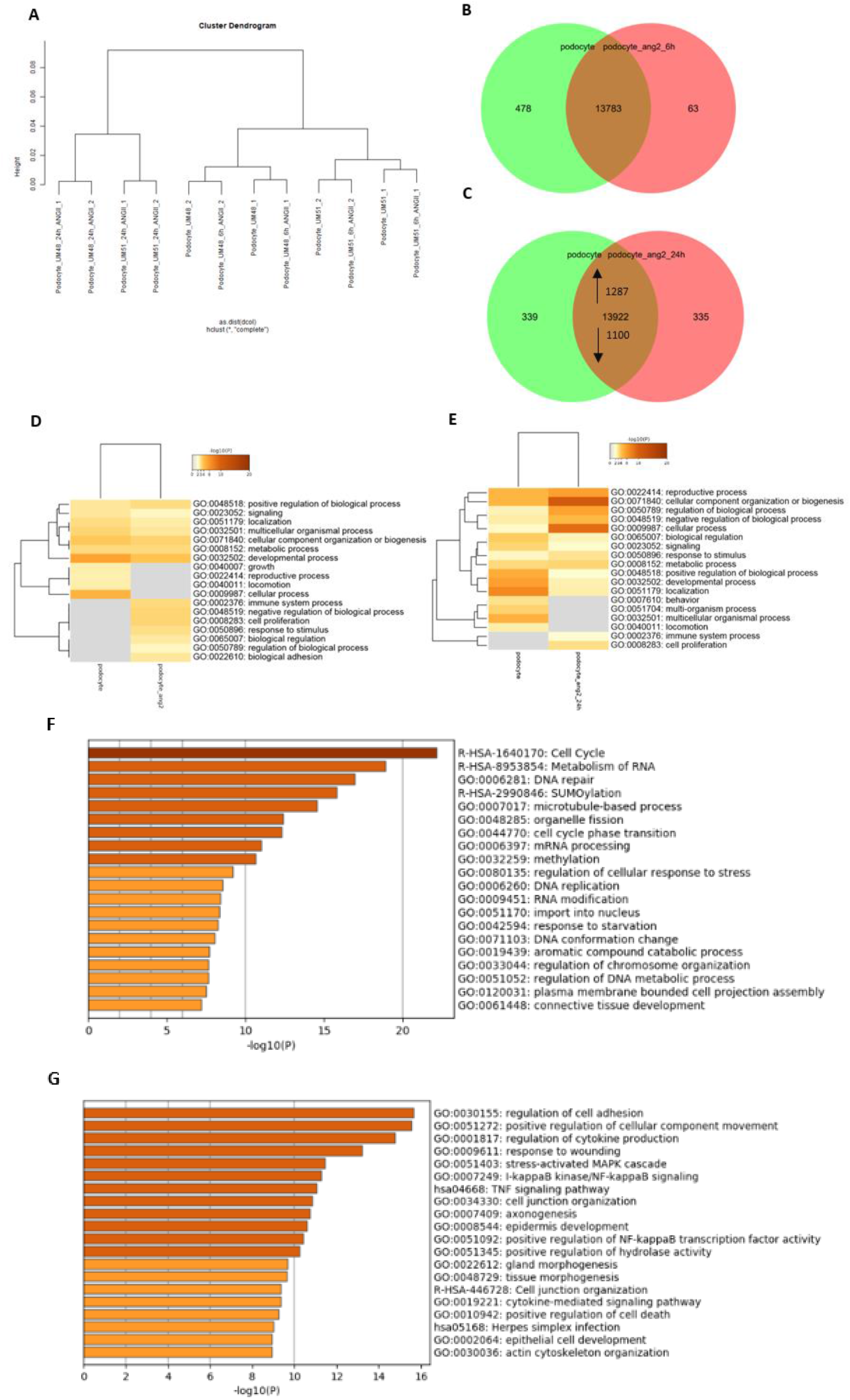
Comparative transcriptome and Gene Ontology analysis of untreated podocytes UM48 and UM51 and treated with Angiotensin II for 6h and 24h. Podocytes were treated with ANGII (100 µM) for 6h and 24h. The hierarchical cluster dendrogram revealed 2 distinct clusters-24h treated and 6h treated and untreated cells as a cluster. (a). Expressed genes (det-p < 0.05) in podocytes and after ANGII treatment are compared by Venn diagrams after 6h (b) and 24h (c), revealing unique and overlapping expression patterns. The most over-represented GO BP-terms (13922 genes) common in either condition are shown in d after 6h of treatment and e after 24h. The terms include locomotion, growth, immune system process, response to stimulus biological adhesion, metabolic process, and cell proliferation for the treated podocytes. The 10 most over-represented GO BP-terms in the up (1287) and down (1100)-regulated genes in podocytes treated for 24h with ANGII in comparison to untreated cells are shown in (f). The upregulated genes are associated with cell cycle, DNA repair, cell cycle phase transition, methylation and regulation of cellular response to stress while downregulated genes are associated with regulation of cell adhesion, cell junction organization, axonogenesis, actin cytoskeleton organization and tissue and gland morphogenesis (g).

By analysing the secretomes of UM48 and UM51 podocytes with and without 6h and 24h of ANGII treatment, a plethora of significantly regulated secreted factors, specific for human kidney and associated with renal diseases, were identified (supplementary fig. 4). In accord with the transcriptome data only minor changes after 6h of ANGII treatment could be observed with only three cytokines significantly altered, ADIPOQ, ANPEP and AGER. Major alterations within the secretome were manifested after 24h, resulting in a significant up-regulation of ANPEP, CCN1, IL6, MMP9, SERPINA3 and VEGF and down-regulation of CXCL16, REN, TNFα, TFF3 and PLAU.

## Discussion

The kidney glomerulus or renal corpuscle consists of a glomerular tuft and the Bowman’s capsule. Its major task is the filtration of blood to generate urine and it consists of four distinct cell types endothelial cells, mesangial cells, parietal epithelial cells of Bowman’s capsule and podocytes. In the present manuscript, we present a differentiation protocol for the successful and reproducible differentiation of UdRPCs into podocytes. These podocytes have the typical complex architecture and express podocyte specific proteins Synaptopodin, Nephrin and Podocin, which are necessary for the establishment of either the slit diaphragm or the foot processes. Podocytes are major contributors towards balancing colloidal pressure within the renal system and transport metabolites by endo/exocytosis of Albumin. As means of diagnosis, the quantitative analysis of Albumin in urine is used as a mark of renal cell functionality. The Albumin infiltration pathway partly takes place within podocytes *in vivo* and *in vitro* [34–36]. The comparison of the transcriptomes pertaining to UdRPC differentiated podocytes with previously reported data-sets of iPSC–derived podocytes, biopsy derived human glomeruli, and mouse podocytes [21], revealed distinct clustering of UdRPCs and their differentiated counterparts, whereas most of the genes were found to be upregulated after our differentiation protocol. Furthermore, genes that are exclusively expressed or upregulated (det-p < 0.05) after our differentiation protocol, but are not on the reported list [21], have been shown to be critical for podocyte morphology and/or function. Unique genes from our data that are associated with the filtration barrier or cytoskeleton of podocytes include the genes *CX3CL1* [37], *DNM3* [38], *DPP4* [39] and *NRX1* [40]. Unique genes from our data that are associated with podocyte function or survival include *BDNF* [41], *CXCL16*, [42] *GRIA3* [43], *PDGF-C* [44], *RAB3A* [45] and *VEGF-C* [46]. Members of Semaphorin gene family *SEMA3A* and *SEMA3C* were also significantly upregulated. Both are guidance proteins, expressed during kidney development and regulate kidney vascular patterning, endothelial cell migration, survival, uteric bud branching and podocyte-endothelial crosstalk [47]. In a mouse model of semaphorin3a gain- and loss of function experiments revealed a dose dependent role of Semaphorin3a in podocyte differentiation and establishment of the glomerular filtration barrier [48].

During renal development the glomerulus is established in four stages: the renal vesicle stage, the S-shaped body stage, the capillary loop stage, and the maturing-glomeruli stage [33]. This developmental process is initiated by the metanephric mesenchyme. It induces the ureteric bud epithelium to grow and finally to branch and form the collecting duct system of the kidney. Hereby the mesenchymal cells of the metanephric mesenchyme have to form the polarized epithelial cells of the renal vesicle and therefore undergo a process that is referred to as mesenchymal-to-epithelial transition (MET) [49]. The podocytes in the glomerulus are generated by a population of cells within the pariental epithelium. These cells are thought to migrate into the glomerular tuft and then differentiate into the mesenchyme like podocytes [50, 51]. Therefore, the maturation of podocytes is associated with a physiological epithelial-to-mesenchymal transition [1]. These developmental processes are also reflected in our transcriptome data. Genes that are upregulated in the derived podocytes are annotated with the GO BP-terms such as axon guidance and chemotaxis, thus enabling speculation that this might reflect the migration process during podocyte maturation. In addition, as differentiation into podocytes also involves both MET and EMT, it is not surprising that the upregulated data set consists of numerous critical effectors for both processes, for instance *wnt-1* has been reported to be a major driver of MET [52]. Interestingly the genes *Cav1* and *Dlg1* have been linked to kidney development and renal pathological conditions in mice [49, 53]. Further signs of EMT/MET are the differential expression of cell-cell contact-associated markers and functional changes associated with the conversion between mobile and stationary cells [54]. Confirmation that these changes are induced by our differentiation protocol is reflected in the podocyte upregulated dataset by the annotated GO BP-terms cell adhesion, cell substrate adhesion, as well as the podocyte exclusive expression of the collagens *COL12A1* and *COL14A1*.

Developmental processes are regulated by epigenetic mechanisms and these are of fundamental importance for cellular differentiation [55]. Epigenetic remodeling is associated with the establishment and removal of histone modifications and DNA-methylation, to generate the cell type-specific epigenome. The epigenetic remodeling needed for the differentiation of UdRPCs into podocytes is also reflected by our transcriptome data. We observed downregulation of the de novo DNA-methyltransferase 3a (*DNMT3a*) as well as Tet Methylcytosine Dioxygenase 2 (*TET2*) in podocytes. Whilst DNMT3a is required for the establishment of CpG methylation during embryogenesis [56], the members of the TET family are needed for the initiation of demethylation [57, 58]. This targeted epigenetic remodeling during the differentiation process is manifested at the *WT1* promoter, as observed by bisulfite sequencing, therefore lending support to the speculation that this process is mediated by TET2. Furthermore, genes associated with to the polycomb repressive complex, *EZH1* and *SUZ12 [59]*, were also found to be downregulated. Specifically, EZH1 has been reported to be important for the maintenance of stem cell identity and execution of pluripotency [60]. Podocytes are terminally differentiated cell, which are unable to undergo cell division in vivo [1]. Due to the observed downregulation of genes associated with epigenetic modifications and cell cycle processes, we propose that the podocytes generated with our protocol have acquired this terminally differentiated state. This hypothesis is further strengthened by the observed downregulation of MSC associated cell cycle related genes *BUBR1, CDC7, CDC25A, MAD2L1, FGFR2, SIX3* and *TOPA2*. Furthermore, these genes have been reported to maintain stemness in mesenchymal stem cells and their downregulation is linked to the differentiation of MSCs [61–68].

Of note the derived podocytes were found to have upregulated the mRNA expression of several solute carrier family members (SLC) (supplementary fig. 3). To our knowledge this is the first report describing the expression of SLCs in human urine-derived podocytes. We propose that the abundance of solute carrier family members is due to the increase in transmembrane transport, as indicated by our podocyte exclusive GO BP-terms, regulation of ion transmembrane transport, neurotransmitter transport, organic acid transport and chemical synaptic transmission. Furthermore, SLCs have been shown to be involved in human diseases and drug targets, thus establishing our human urine-derived podocytes as a valuable cellular tool for studying SLCs in renal development and under pathological conditions [69].

The renin-angiotensin system (RAS) major implications in humans is the maintenance of plasma sodium concentration, arterial blood pressure and extracellular volume. Activation of the RAS leads to hypertension, cell proliferation, inflammation, and fibrosis and therefore has implications for all tissues of the body [70]. Angiotensin receptors and especially AGTR1 in the kidney have been reported to be primarily causative for hypertension in mammals [71] and are associated with renal sodium handling. AGTRs therefore have been recognized to be commonly expressed in several segments of the nephron including the thick ascending limb, distal tubule, collecting duct and renal vasculature [72–77]. The stimulation of these receptors has been linked to renal vasoconstriction, reduced medullary blood flow, diminishing the renal capacity of sodium handling [78–80], leading to podocyte injury and loss [15]. The main mediator of RAS is ANGII and it has been shown that ANG receptors within the kidney are the main mediators of hypertension [71]. Elevated levels of circulating and intracellular ANGII and aldosterone lead to pro-fibrotic, -inflammatory and - hypertrophic milieu that causes remodelling and dysfunction in cardiovascular and renal tissues [81]. The major finding in this study is the disruptive effect of ANGII on the cytoskeleton of podocyte. Since ANGII obviously interferes with the podocyte cytoskeleton, it is important to note that commonly housekeeping genes like *ß-ACTIN* are unsuitable for normalization and we therefore used *RPL0*, which should not be affected. Employing immunofluorescence-based protein expression detection we observed cytoskeletal changes of the normal morphology of podocytes, leading to the inhibition of podocyte spreading and down-regulated expression of *NPHS1* and *SYNPO*. The down-regulated expression of NPHS1 and SYNPO has been reported as a hallmark of podocyte injury [13] and is causative for the disruption of foot processes (FPs) and the slit diaphragm (SD). This effect also probably due to the downregulated expression of numerous genes associated with the GO-BP-actin cytoskeleton organization (fig. 5g and supplementary table 9). As an example instance Alpha-Actinin-4 (ACTN4) is highly expressed in podocyte FPs and NPHS1 is connected to the actin cytoskeleton thus contributing to podocyte actin dynamics, signaling and mobility [2, 3, 6]. Furthermore, our transcriptome data unveiled the downregulated expression of *CORO2B, DPP4, ELF3, LDLR, KLF4, ROBO1* and *WNT7A*. These genes have been linked to podocyte injury and filtration barrier impairments [18, 39, 82–86]. Of note, *KLF4* has been shown to directly regulate *NPHS1* expression [39], thus suggesting that the disruptive effect of ANGII might be amongst others also mediated by *KLF4*. In the upregulated gene set numerous genes associated with cell cycle initiation and DNA double strand repair, such as *FANCD2, MSH2, WDR7* and *XRCC4* (fig. 5f and supplementary table 8). In a mouse model of Brand et al., it was shown that ANGII is indeed able to induce DNA damage via a dose dependent increase of oxidative stress, which results in DNA damage [87]. Furthermore, elevated levels of *ATF4, EDN-1, TGFB2* and *PLK2* have all been linked to podocyte injury and loss of key proteins necessary for the maintenance of the complex podocyte architecture [88–91]. Most of the cytoskeleton-related genes that were downregulated upon ANGII treatment in podocytes, had lower levels of expression in the undifferentiated UdRPCs. Although the expression of Ankyrins, Collagens, Keratins and Semaphorins have been linked to kidney development and functionality [38, 47, 48, 92], to our knowledge only Semaphorins have been reported to be involved in kidney injury [47, 93, 94]. Of note numerous members of the sodium and chloride symporter family SLC6 [69], are highly upregulated in the differentiated podocytes (supplementary fig. 3), while upon ANGII stimulation upregulated expression of AGTR1 and 2 was observed. We conclude that our findings offer the possibility of establishing biomarkers indicative of differentiation of UdRPCs into podocytes as well as initiation and progression of CKD.

To surmise, we have described a differentiation protocol for the successful and reproducible differentiation of SIX2-positive urine derived renal progenitor cells (UdRPCs) from two African males aged 48 and 51 and one female Caucasian aged 45 years into mature podocytes with the ability to execute Albumin endocytosis. UdRPCs enable the unique possibility to study nephrogenesis and associated diseases thus obviating the need of iPSCs. Furthermore, the responsiveness of UdRPC derived podocytes to ANGII implies potential applications in nephrotoxicity studies and drug screening. Figure 6 presents a graphical summary of this study.

**Figure 6:**
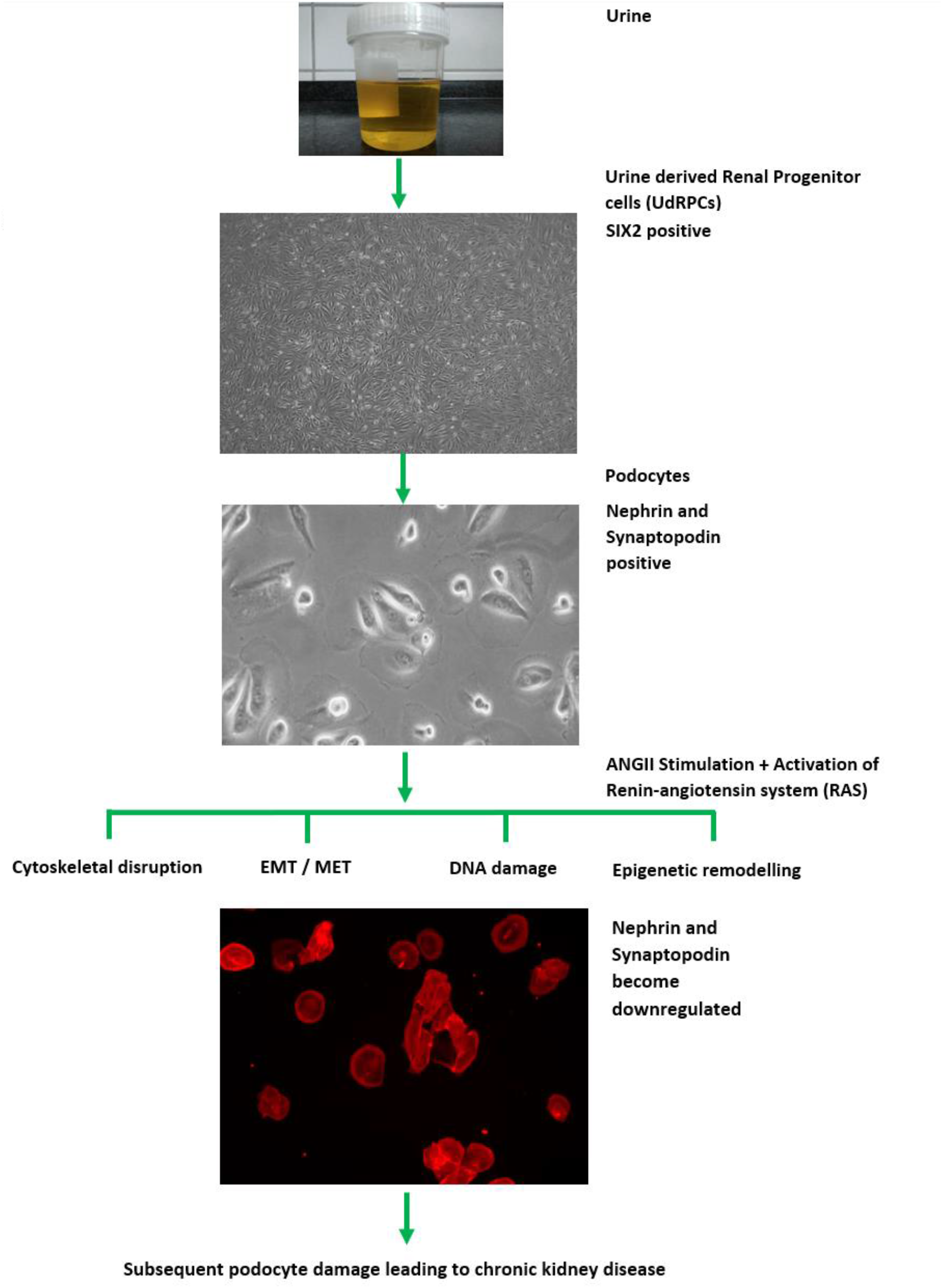
Graphical summary of this study. SIX2-positive renal progenitor cells were isolated directly from human urine and cultured in proliferation medium. UdRPCs differentiated into podocytes, by low density cultivation in advanced RPMI medium supplemented with 30 µM retinoic acid. Stimulation with 100 µM ANGII results in cytoskeletal remodelling. Furthermore, the transcriptome revealed altered expression of genes associated with epigenetic remodelling, DNA damage response, EMT and MET. ANGII activated the renin-angiotensin system resulting in podocyte damage, -loss and ultimately leading to chronic kidney disease.

## Supporting information

supplementary table 4 UdRPC vs podocyte

supplementary table 5 Podocyte up

supplementary table 6 Podocyte down

supplementary table 7 Podocyte vs 24h angII

supplementary table 8 24h angII up

supplementary table 9 24h angII down

## Acknowledgements

James Adjaye acknowledges funding from the medical faculty of Heinrich Heine University-Duesseldorf, Germany

## Supporting information

Supporting information in the supplementary section or from the author.

