## supplementary figures for "Angiotensin II disrupts the cytoskeletal architecture of human urine-derived podocytes and results in activation of the renin-angiotensin system"

Table 1: RT-qPCR Primers

| **Primer name** | **Sequence** | **Annealing temperature (°C)** | **Product length**  **(bp)** |
| --- | --- | --- | --- |
| AGTR1 s | 5’- tct cag cat tga tcg ata cc -3’ | 60 | 80 |
| AGTR1 as | 5’- tga ctt tgg cta caa gca tt -3’ |  |  |
| AGTR2 s | 5’- tat ggc ctg ttt gtc ctc at -3’ | 60 | 114 |
| AGTR2 as | 5’- cat tgg gca tat ttc tca gg -3’ |  |  |
| LMX1b s | 5’- cat cct cac cac gca gca g -3’ | 60 | 155 |
| LMX1b as | 5’- tct tca tct ttg ctc ttt ggt tct -3’ |  |  |
| NPHS1 s | 5’- gcg ggt tct gct acg atg gtg -3‘ | 60 | 295 |
| NPHS1 as | 5‘- caa aca cac cag cct cac ccg -3’ |  |  |
| RPL s | 5’- tcg aca atg gca gca tct ac -3’ | 60 | 195 |
| RPL as | 5’- atc cgt ctc cac aga caa gg -3’ |  |  |
| SYNPO s | 5-‘ ccc caa cct ctc ctc taa cc -3’ | 60 | 116 |
| SYNPO as | 5’- atg aca cag gag gca gaa gaa t -3‘ |  |  |
| WT1 s | 5’- cac agc aca ggg tac gag a -3’ | 60 | 133 |
| WT1 as | 5’- caa gag tcg ggg cta ctc c -3’ |  |  |

Table 2: bisulfite sequencing primers

| **Primer name** | **Sequence** | **Annealing temperature (°C)** | **Product length**  **(bp)** |
| --- | --- | --- | --- |
| NPHS2 bseq s | 5’- gtt ttg agg atg gag agg agg -3’ | 55 | 415 |
| NPHS2 bseq as | 5’- ttc cta aaa acc taa aca tcc aac -3’ |  |  |
| WT1 bseq s | 5’- ggg gga ggg ttg tgt tat at -3’ | 54 | 270 |
| WT1 bseq as | 5’- ctc ctt acc cca acta cc taa cta c -3’ |  |  |

Table3: antibodies

| **Antigen** | **Company** | **Dilution** |
| --- | --- | --- |
| α-ACTININ | Abcam (#108198) | 1:200 |
| CD2AP | Cell Signaling Technology (A599) | 1:200 |
| NPHS1 | Invitrogen (#PA5-20330) | 1:200 |
| NPHS2 | Abcam (#50339) | 1:200 |

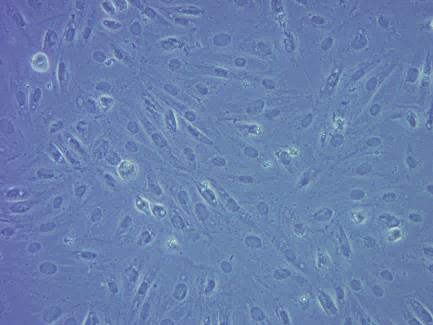

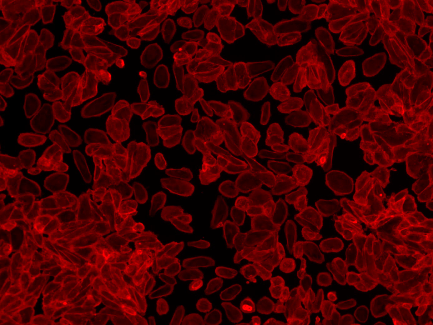

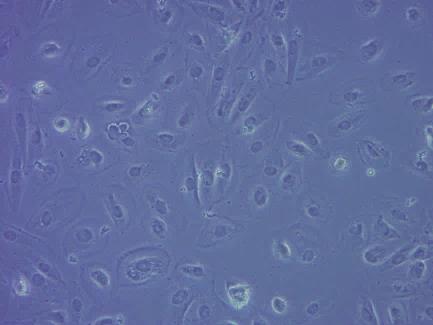

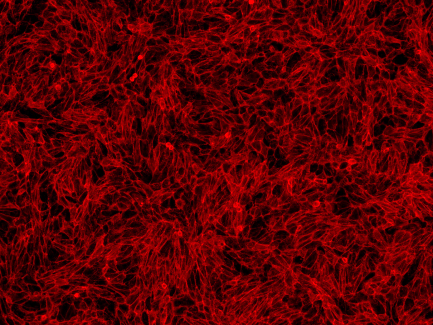

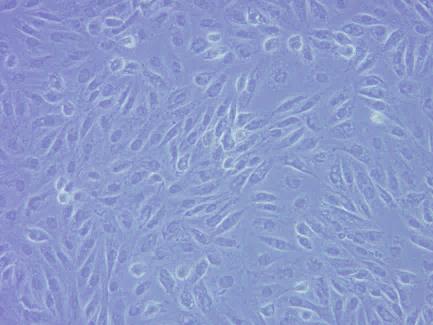

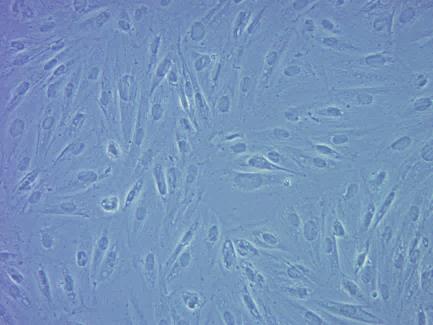

**A**

**Control high density**

**UM 51**

**UM 48**
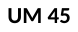

**UM 45**

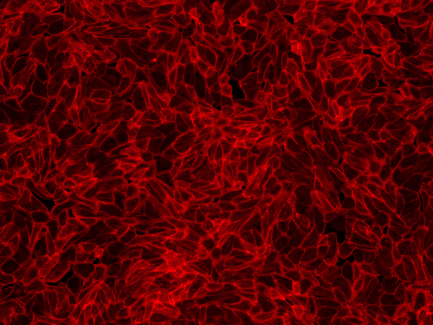

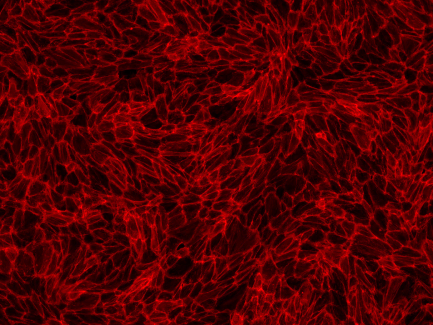

**Control high density**

**100µM ANGII high density**

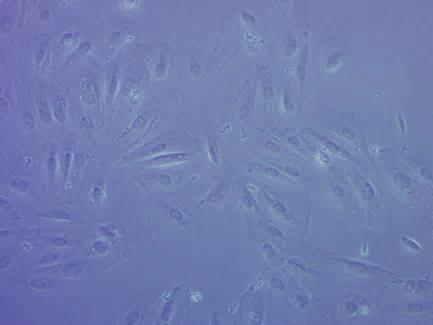

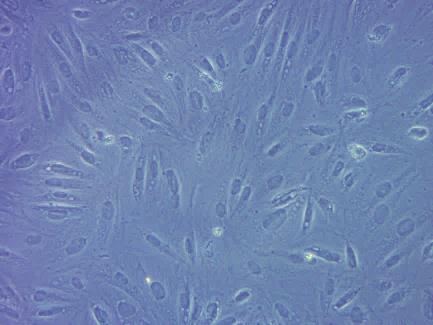

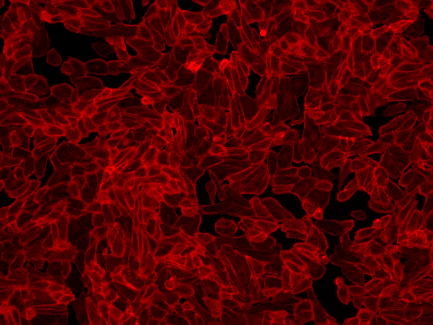

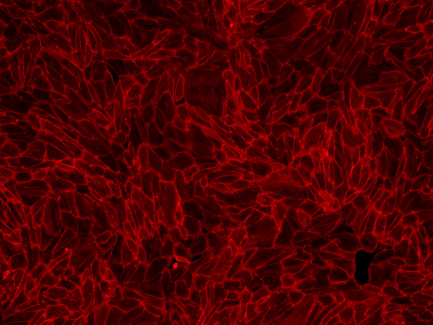

**100µM ANGII high density**

**C**

**B**

|  | AGTR2 expression |
| --- | --- |
| Podocyte UF45 | 2.0037 |
| Podocyte UF45 6h ANGII | 1.40932 |
| Podocyte UM48 | 2.15397 |
| Podocyte UM48 6h ANGII | 2.50649 |
| Podocyte UM51 | 1.22123 |
| Podocyte UM51 6h ANGII | 7.11732 |

|  | AGTR1 expression |
| --- | --- |
| Podocyte UF45 | 1.41144 |
| Podocyte UF45 6h ANGII | 2.2338 |
| Podocyte UM48 | 1.3904 |
| Podocyte UM48 6h ANGII | 1.87276 |
| Podocyte UM51 | 1.7409 |
| Podocyte UM51 6h ANGII | 1.99423 |

**E**

**D**

|  | SYNPO expression |
| --- | --- |
| Podocyte UF45 | 1.04006 |
| Podocyte UF45 6h ANGII | 1.3632 |
| Podocyte UM48 | 1.92919 |
| Podocyte UM48 6h ANGII | 1.09936 |
| Podocyte UM51 | 4.06945 |
| Podocyte UM51 6h ANGII | 1.37777 |

|  | NPHS1 expression |
| --- | --- |
| Podocyte UF45 | 1.3293 |
| Podocyte UF45 6h ANGII | 0.824733 |
| Podocyte UM48 | 2.30537 |
| Podocyte UM48 6h ANGII | 0.76693 |
| Podocyte UM51 | 2.94105 |
| Podocyte UM51 6h ANGII | 0.417206 |

**Supplementary figure 1: Dynamic changes in the morphology of human urine derived podocytes after treatment with Angiotensin II (ANGII) after 6h.**

Human urine derived renal progenitor cells, derived from three different individuals, have been differentiated into podocytes, by high density cultivation in advanced RPMI medium supplemented with 30 µM retinoic acid. The top panel displays the typical „fried egg“ shaped podocyte morphology. The lower two panels show the morphology changes after 6h of 100 µM ANGII treatment. Podocyte cytoskeleton was visualized by immunofluorescent staining’s for α –ACTININ in red (a).

ANG II interferes with the cytoskeleton of the podocytes, inhibiting podocyte spreading and resulting in the loss of foot processes and the observed roundish phenotype (indicated by the white arrows). Expression of podocyte transcription factors AGTR1 (b), AGTR2 (c), *NPHS1* (d) and *SYNPO* (e) were determined by quantitative real time PCR normalized by RPL0.

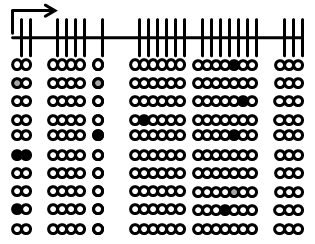

**Podocyte UM51**

***NPHS2***

***NPHS2***

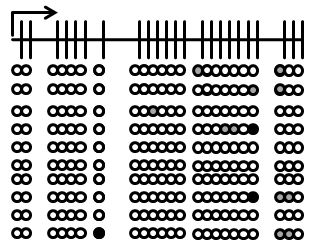

**Podocyte UM51**

**+ 6h ANGII**

**Supplementary figure 2: DNA Methylation changes at the NPHS1 promoter upon 6h of ANGII treatment.**

Human urine derived renal progenitor cells, derived from three different individuals, have been differentiated into podocytes, by low density cultivation in advanced RPMI medium supplemented with 30 µM retinoic acid. Podocyte injury was induced by adding 100 µM ANGII for 6h. Bisulfite genomic sequencing of a 415bp long NPHS2 fragment slightly upstream of the TSS, spanning 23 CpG-dinucleotides, give detailed information about the DNA Methylation changes occurring during podocyte injury. Black, white and grey circles stand for methylated, unmethylated and undefined CpG dinucleotides, respectively.

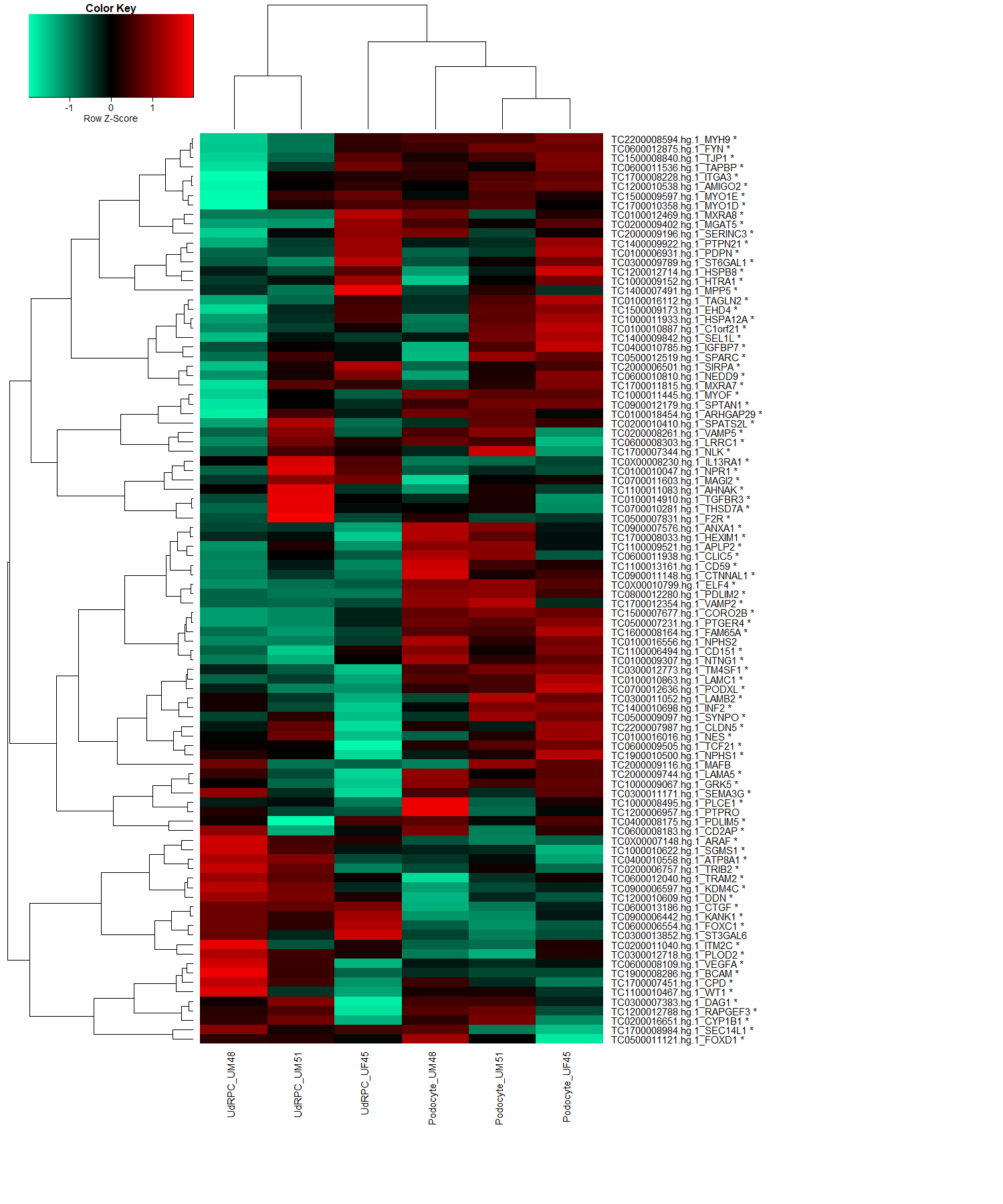

**Supplementary figure 3: iPS cell–derived podocytes, kidney biopsy isolated human glomeruli, and mouse podocytes and Solute Carrier (SLC) Family members, reveals distinct expression patterns of UdRPCs and their differentiated counterparts.**

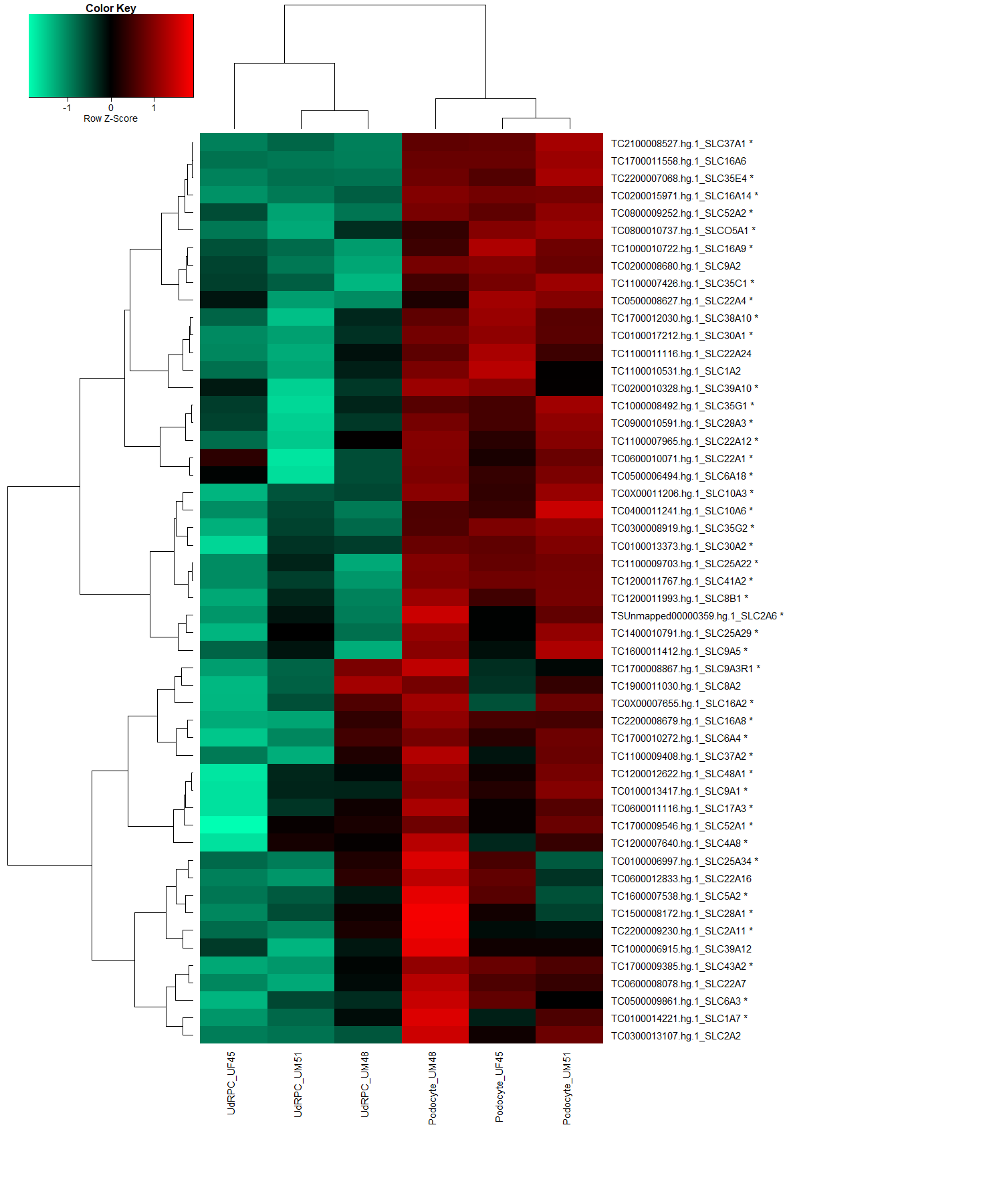

**Supplementary figure 3: continuing.**

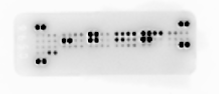

UM48

UM51

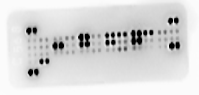

control

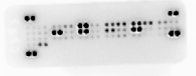

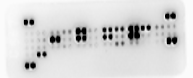

6h ANGII

(100 µM)

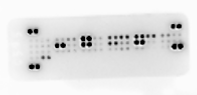

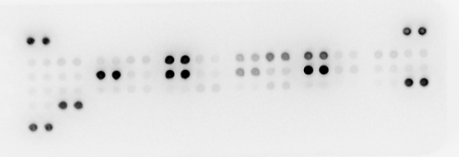

24h ANGII

(100 µM)

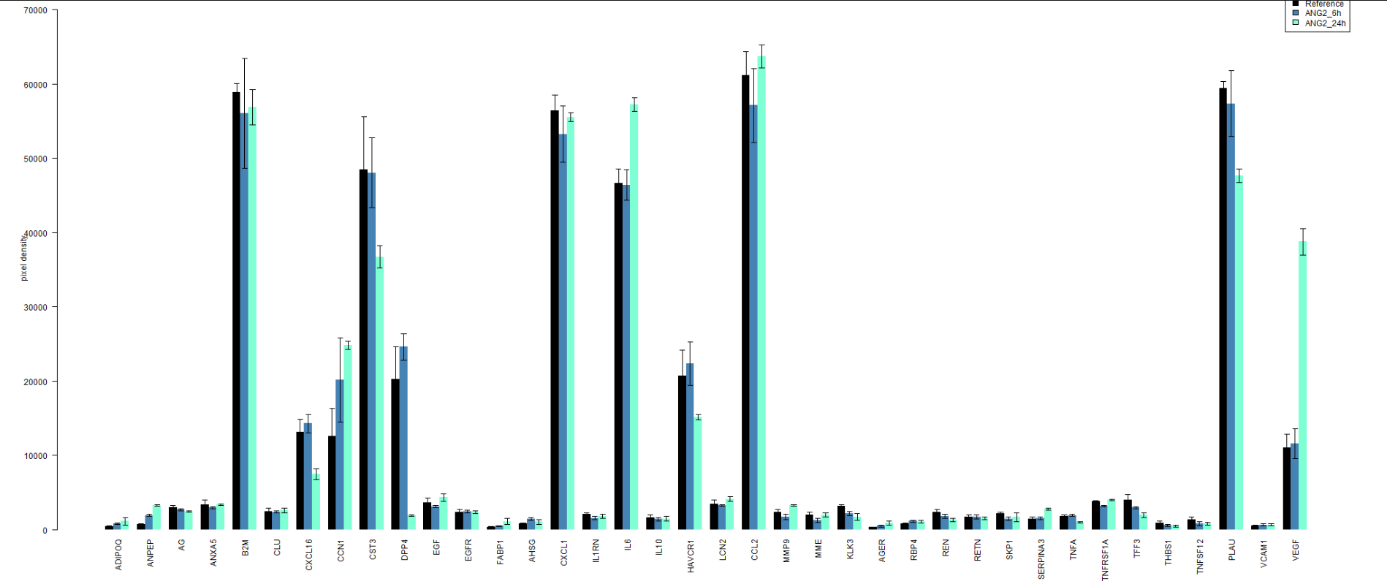

**Supplementary figure 4: Secretome membranes and analysis of podocytes with and without 6h and 24h of Angiotensin II treatment.**
